# 1-Methyl-tryptophan enantiomers differently affect the cross-talking between mononuclear and tumor cells and interferon-γ production

**DOI:** 10.1101/578369

**Authors:** Maryana Stephany Ferreira Branquinho, Maysa Braga Barros Silva, Renan Orsati Clara, Ariane Rivellis Julio, Silvya Stuchi Maria Engler, Ernani Pinto Junior, Ana Campa

## Abstract

The inhibition of the enzyme indoleamine-2,3-dioxygenase (IDO), that catalyzes the oxidation of the amino acid tryptophan to kynurenine (KYN), is considered a good target for immunoadjuvants in antineoplastic therapy. 1-Methyl-tryptophan (1-MT) is the most studied molecule for this purpose. Although L-1-MT is better than D-1-MT in inhibiting IDO, for an unknown reason the D-enantiomer has higher clinical efficacy. Here we took advantage of co-cultures of tumor cells (SK-Mel 19 melanoma line; 1×10^5^ cells/well) with peripheral blood mononuclear cells (PBMC; 5×10^6^ cells/well) to verify the effect of 1-MT enantiomers on cytokine production and tumoricidal activity. At a concentration that did not affect KYN production, 1-MT (50 µM) affected the production of TNF-α, IL-10, and IFN-γ measured in co-cultures supernatants. Stereospecificity was only observed for IFN-γ production. D-1-MT inhibited more than 30% of IFN-γ production, while L-1-MT had no effect. Stereospecific effect was also seen in PBMC tumoricidal activity, estimated by tumor cell viability (Trypan assay). The racemic mixture DL- and D-1-MT almost doubled the tumoricidal activity of PBMCs, while L-1-MT had no effect. These are previous unknown off-target effects of D-1-MT. Our data suggest the modulation of IFN-γ and the activation of tumor recognition and killing processes by immune cells as important features for the in vivo effects of the D-1-MT. These findings should be considered in future studies of immunoadjuvants for cancer treatment.

## Introduction

A main approach given to cancer treatment is a therapeutic regimen that combines classical antineoplasic agents with immunoadjuvants. Among the available options of immunoadjuvants, the inhibition of the enzyme indoleamine-2,3-dioxygenase (IDO) seems to be one of the most promising therapy strategies to prevent tumor escape (Zhai LJ, *et al*., 2015; Prendergast GC, *et al.*, 2014). In various types of cancer, IDO is upregulated and is associated with poor prognosis (Burugu S, *et al*., 2017; Selvan, Senthamil R, *et al*., 2016).

IDO is an immunoregulatory enzyme that catalyzed the rate-limiting step of tryptophan (Trp) metabolism through the kynurenine (KYN) pathway. T-cells and antigen-presenting cells (APCs) respond to Trp depletion and KYN production promoting a tolerogenic environment (Platten M, *et al*., 2015). Cell-cycle arrest and anergy of effector T-lymphocytes caused by changes in Trp metabolism results in immune tolerance. Moreover, the accumulation of KYN induces T-cells apoptosis and, in T naïve cells, the binding of KYN to the aryl hydrocarbon receptor (AhR) generates a T regulatory cells (Treg) phenotype (Zhai LJ, *et al*., 2015; Becker JC, *et al.,* 2013). IDO inhibition has been proposed as an adjuvant improving immunosurveillance in a dozen of ongoing clinical trials for antincancer therapy. Despite growing interest in the search for new IDO inhibitors, the classical IDO inhibitor 1-methyl tryptophan (1-MT) has the molecular characteristics for a good inhibitor and is the chief molecule screened in the majority of the clinical studies.

1-MT is a competitive inhibitor of IDO that exists in the enantiomeric forms L-methyl-tryptophan (L-1-MT) and D-methyl-tryptophan (D-1-MT). When recombinant human IDO enzyme is tested in a cell-free assay system, L-1-MT is more efficient than D-1-MT in inhibiting both IDO enzyme isoforms (IDO1 and IDO2) (Hou DY, *et al*., 2007). The same was observed for IDO in cell cultures (Qian F, *et al.,* 2013). L-1-MT has a *Ki* value of 19 µmol/L, whereas D-1-MT has a *Ki* of ~100 µmol/L and the racemic mixture has a *Ki*_DL_ of 35 µmol/L (Hou DY, *et al*. 2007). In spite of the *Ki* values, D-1-MT show higher antitumor activity in vivo in different tumors and combined therapy regimens. D-1-MT was more effective in reducing tumor mass in experimental models (Hou DY, *et al.,* 2007; Liu K T, *et al.,* 2016), and for that reason it is used in Phase 1 and 2 clinical trials with the commercial name of *Indoximod*^®^ (Tang SC, *et al.,* 2016). Indoximod has been trial in mono and combined therapy to treat sarcoma, NSCLC, melanoma and colorectal cancer (Yentz S and Smith D, 2018). Although the initial focus on 1-MT took into account its property to inhibit IDO, currently it is clear that many of 1-MT effects in vivo are due to its off-targets effects (Fox E *et al*., 2018). The reasons why D-enantiomer has a higher efficiency are not clear and possible causes are related to stereoselective interaction with optically active biological enzymes, bioavailability, pharmacokinetics and other off-target effects.

Herein we went further looking for the different effects of D-1-MT and L-1-MT on the interaction between tumor and immune system including the production of some cytokines important in cancer. The cross-talking between tumor and immune cells was determined by the cytotoxic activity of peripheral blood mononuclear cells (PBMC) against the melanoma lineage SK-Mel 19.

## Materials and Methods

### PBMC isolation

Human peripheral blood mononuclear cells (PBMCs) were isolated from peripheral venous blood obtained from healthy volunteers by centrifugation over a Ficoll– Hypaque gradient (Histopaque; d=1.077) (Boyum A, 1968). Cell concentration and viability were determined in a Neubauer chamber. This study was approved by the local research ethics committee (Comitê de Ética em Pesquisa-FCF/USP, no. 1.862.199 -December 12, 2016).

### Tumor-PBMC Co-culture

Melanomas (SK-Mel 19 - 1×10^5^ cells/well) and PBMCs (5×10^6^ cells/well) were co-cultured in a twelve-well plate in RPMI-1640 (GibcoW) in the presence of DL-1-MT, D-1-MT and L-1-MT (50µM). The ratio melanoma:PBMC of 1:50 was the small ratio tested able to cause a significant death of tumor cell. After 72 h, tumor cells were counted with Trypan blue and the supernatant was used for quantification of cytokines by Cytometric Bead Array (CBA) and of KYN by LC-ESI-MS/MS.

### Cytokine determination

The concentration of TNF-α, IL-10, IFN-γ, IL-8, IL-1β, IL-12 and IL-6 was simultaneously analyzed by flow cytometry (FACS Canto II; Becton-Dickinson) using the BD Cytometric Bead Array (CBA) Human Inflammatory Cytokines Kit (BD™) according recommendations. For the analysis we used 25 μL of the supernatant from the co-cultures. The data were analyzed using FCAP Array ^TM^ Software.

### KYN quantification

The KYN supernatant concentration was analyzed for LC-ESI-MS/MS (LC 1250 Bin Pump VL and 1260 HiP ALS autosampler coupled to a triple quadrupole 6460 mass spectrometer - all from Agilent Technologies, CA, USA). Electrospray ion source (ESI) was operated in the positive mode. MS data were acquired by multiple reaction monitoring (MRM) mode and analyzed by MassHunter Quantitative Analysis from Agilent Technologies. Phenomenex (Torrance, CA, USA). Luna C18 (2) (150 mm x 2 mm, 3 μm) reversed-phase column was used for LC separation. Chromatography was performed by a gradient elution composed by ammonium formate 10 mM in water (adjusted to pH 2 with formic acid) (A) and acetonitrile (B). Injection volume was 5 μL applied at a flow rate of 0.2 mL.min^-1^. A standard stock solution was prepared in MeOH (from 0.01 μM to 50 μM), and the linearity of the dependence of response on concentration was verified by regression analysis.

### Statistics

The statistical significance of differences in the mean values of all experimental groups was calculated using a ONE Way ANOVA. P values < 0.05 were considered to be statistically significant.

## Results

To conduct this study, we choose a 1-MT concentration that did not inhibit KYN synthesis. This precaution was taken to identify additional effects of 1-MT beyond IDO inhibition. Usually, 1mM of 1-MT is used at to inhibit IDO in vitro (Moreno ACR, *et al*., 2013). We choose 50 μM 1-MT that was a concentration insufficient to inhibit KYN production. In our co-culture condition the concentration of KYN was approximately 25 μM and the addition of 1-MT enantiomers did not caused changes in this concentration **(Figure 1A)**.

**Figure 1:**
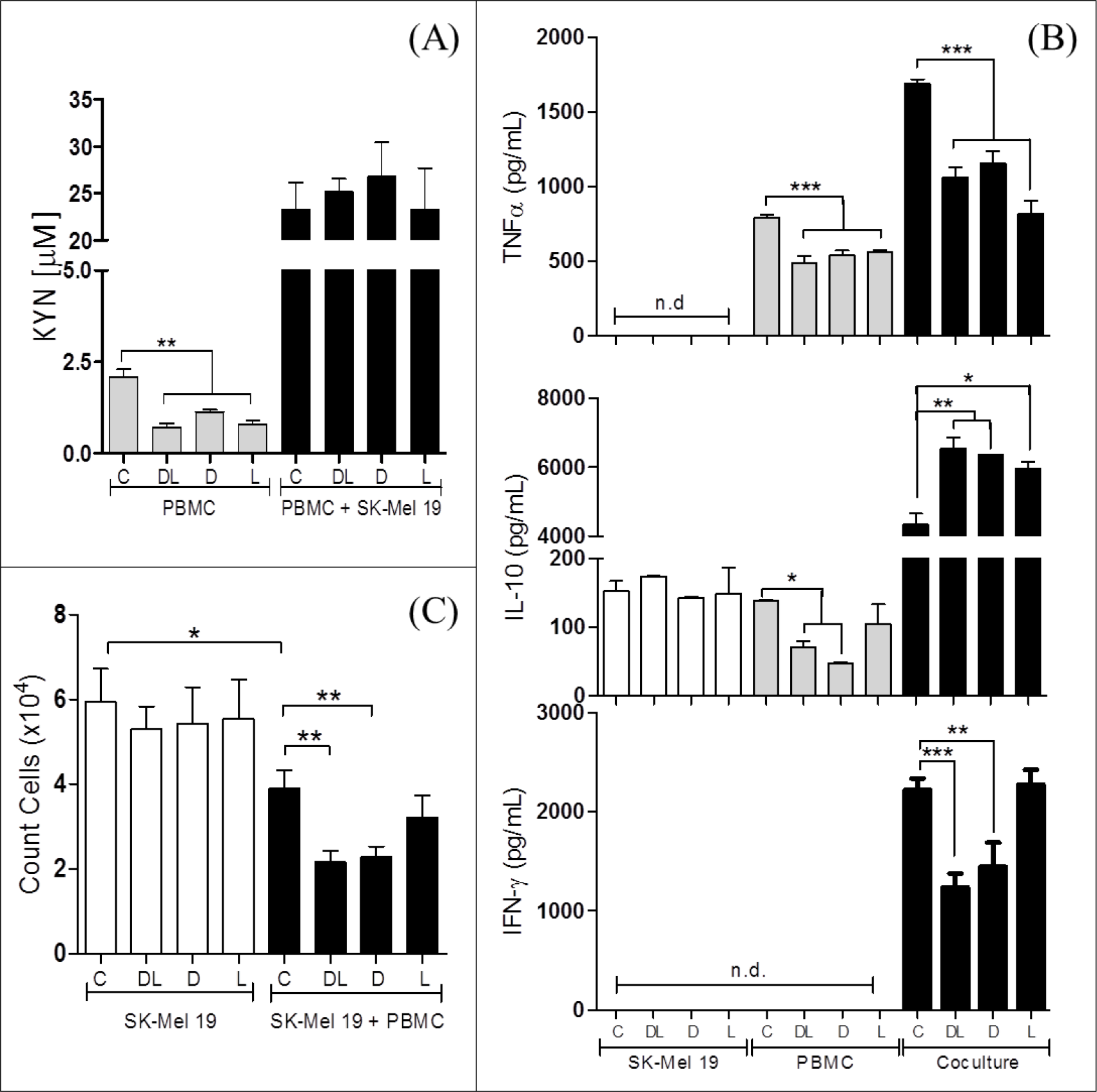
D-1-MT inhibits IFN-γ production and enhances the tumoricidal activity of PBMCs. The production of kynurenine (KYN) by PBMC (5×10^6^ per well) and SK-Mel 19 (1×10^5^ per well) co-cultures was not affected by D-, L- and DL-1-MT enantiomers (50 µM) **(A)**. At these conditions, 1-MT affected TNF-α, IL-10, and IFN-γ production measured in co-cultures supernatants (**B**). Stereospecificity was only observed for IFN-γ production. D-1-MT inhibits more than 30% of IFN-γ production. Stereospecific effect was also seen in PBMC tumoricidal activity, estimated by tumor cell viability (Trypan assay). D- and DL-1-MT almost double the tumoricidal activity of PBMCs (**C**). n=7, 72h co-cultures. * p <0.05, **p <0.01, ***p <0.001 for 72 h.

To evaluate the effect of 1-MT on cytokine production, we measured cytokines in the supernatant of melanoma, PBMC and melanoma:PBMC co-cultures. We measured a set of cytokines known to affect tumor growth, such as TNF-α, IL-10, IFN-γ, IL-8, IL-1β, IL-12 and IL-6. The contact of immune with tumor cells usually leads to incremental production of cytokines (Jehs T, *et al.,* 2014) and we observed this increase for most of the cytokines measured (data not shown) but the effect of 1-MT on cytokine production was only observed for TNF-α, IL-10, IFN-γ (**Figure 1B**). TNF-α produced by PBMC and by melanoma:PBMC was inhibited by 1-MT (~30% and 40%, respectively). IL-10 was also affected by 1-MT. In this case, produced IL-10 was increased by approx. 40% in co-culture. However, we did not observe any stereospecific response for TNF-α and IL-10, there is no differences between D- and L-1-MT at all. Among the cytokines, the most notable difference was observed for IFN-γ; an inhibitory effect of approx. 40% was observed for D-1-MT while no effect at all was observed for the L-enantiomer.

We also use the same conditions of co-cultures as above to access the cross talking between tumor and immune cells leading to tumoricidal activity. At the concentration of 50 μM none of the enantiomers of 1-MT affected tumor cell viability (**Figure 1C, white bars**). The ratio tumor: PBMC used was 1:50, and we observed a reduction of approx. 35% in the number of tumor cells (comparison between the controls). Although 1-MT did not cause any direct effect on tumor cell viability, D-1-MT contributed to a more effective tumor-reactive response by the PBMCs (**Figure 1C, black bars**). This effect was stereospecific and restricted to D-1-MT and to the racemic mixture DL-1-MT.

## Discussion

This study expanded the understanding of the off-targets effects of 1-MT (Moreno ACR, *et al*., 2013; Fox E, *et al.,* 2018), showing stereospecific effects of the D-1-MT enantiomer. In a concentration that precluded IDO inhibition, D-1-MT lead to changes in tumor-reactive response of PBMC increasing the capability of PBMC to induce death of tumor cells and decreased the production of IFN-γ.

Understanding how 1-MT affects the immune response is essential in guiding attempts to optimize its therapeutic efficacy in infections and cancer. 1-MT has already been tested in co-culture of T-lymphocytes and dendritic cells on a viral infection model (Ajamian F, *et al.,* 2015), but not in co-cultures of tumor and immune cells. PBMC provide different populations of lymphocytes and antigen-presenting cells (CD4^+^, CD8^+^, Τ cells, CD4^+^Tregs, CD8^+^Tregs, Th17 cells, monocytes, myeloid DCs, and plasmacytoid DCs) and has been used to mimic in vivo responses (Voo KS, *et al*,. 2014). Here we used co-cultures of tumor cells with PBMC as an easy assay to mimic tumor infiltrating immune cells. Tumor-infiltrating lymphocytes and monocyte-derived cells are observed in several types of cancers and depending on their phenotype pro-or anti-tumor effects are expected (Galdiero *et al*., 2012; Badalamenti *et al*., 2018). By the contact with tumor cell, mononuclear cells can be activated and can cause tumor death or arrest that was used to estimate how 1-MT modulate the immune response and affect the tumoricidal activity.

Given that we used a 1-MT concentration that did not cause modification in the KYN concentration found in the co-culture **(Figure 1A),** we can exclude IDO inhibition in the selective response observed for D-1-MT **(Figure 1A and 1B).** The mechanisms for the increment in the cytotoxic effect of PBMC induced by D-1-MT were not explored in this study and may include different steps from recognition to activation of the immune system. It is interesting to consider that our finding may be associated with the signaling induced by D-amino acids on immune system. Bacteria synthesize D-amino acids (Radkov AD, *et al.,* 2014) and the immune system uses the presence of D-amino acids to sensing bacteria (Fura JM, *et al*., 2014). Could D-1-MT signal the recognition and activation of the immune system contributing to its antitumor effects?

Another interesting point is to consider that the antitumor effect of D-1-MT in vivo may be due to IFN-γ suppression. Although there are many studies showing the role of IFN-γ in the antitumor immune response, opposite effects has been continuously emphasized (Zaidi MR, *et al.,* 2011; Mojic M, *et al.,* 2017; Castro F, *et al*., 2018). In the past IFN-γ was used in clinical trials, however the majority of these trials fail to present conclusive or satisfactory results (Schiller JH, *et al*., 1996). Depending on the doses, IFN-γ treatment was shown to induce resistance of tumor cells to NK cells. IFN-γ was also shown to induce proliferation of tumor cells and is associated to a more aggressive phenotype of tumor cells and metastatic markers (Brocker EB, *et al.,* 1988; Lollini PL, *et al.,* 1993).

In conclusion, we showed previously unknown off-target effects of D-1-MT on the cross talking between tumor and immune cells. Enantiomeric specificity was observed for IFN-γ production and antitumor effect of immune cells. Differences in pharmacokinetics, bioavailability and safety was used to explain the different responsiveness to the treatment between D- and L-1-MT enantiomers (Garbe C, *et al*., 1990), but it seems clear from our results that a broader spectrum of characteristics plays a role in enantiomers specificity, suggesting a central role of IFN-γ modulation and immune response.

## Acknowledgment

This study was supported by Fundação de Amparo a Pesquisa do Estado de São Paulo – FAPESP (process 201509618-2), the Conselho Nacional de Desenvolvimento Científico e Tecnológico – CNPq (process 471510/2010-6), and the Coordenação de Aperfeiçoamento de Pessoal de Nível Superior – Capes. The authors declare no conflict of interest.

## References

Ajamian F, Wu Y, Ebeling C, Ilarraza R, Odemuyiwa SO, Moqbel R, et al. (2015). Respiratory syncytial virus induces indoleamine 2,3-dioxygenase activity: a potential novel role in the development of allergic disease. Clinical and Experimental Allergy. 45(3):644–59 doi 10.1111/cea.12498.

Badalamenti G, Fanale D, Incorvaia L, Barraco N, Listì A, Maragliano R, et al. (2018). Role of tumor-infiltrating lymphocytes in patients with solid tumors: Can a drop dig a stone? Cellular Immunology. doi: 10.1016/j.cellimm.2018.01.013

Becker JC, Andersen MH, Schrama D, Thor Straten P. (2013). Immune-suppressive properties of the tumor microenvironment. Cancer Immunology Immunotherapy. 62(7):1137–48 doi: 10.1007/s00262-013-1434-6.

Boyum. (1968). Isolation of mononuclear cells and granulocytes from human blood. Isolation of monuclear cells by one centrifugation, and of granulocytes by combining centrifugation and sedimentation at 1 g. Scand. J. Clin. Lab. Invest. 21 77–89.

Brocker EB, Zwadlo G, Holzmann B, Macher E, Sorg C. (1988). Inflammatory cell infiltrates in human-melanoma at different stages of tumor progression. International Journal of Cancer. 41(4):562–7 doi: 10.1002/ijc.2910410415.

Burugu, S, Dancsok, AR, Nielsen, TO. (2018). Emerging targets in cancer immunotherapy. Seminars In Cancer Biology, 52, 39–52. doi: 10.1016/j.semcancer.2017.10.001

Castro F, Cardoso AP, Gonçalves RM, Serre K, Oliveira MJ. (2018). Interferon-Gamma at the Crossroads of Tumor Immune Surveillance or Evasion. Front Immunol. doi: 10.3389/fimmu.2018.00847.

Fox E, Oliver T, Rowe M, Thomas S, Zakharia Y, Gilman PB. (2018). Indoximod: An Immunometabolic Adjuvant That Empowers T Cell Activity in Cancer. Front Oncol. doi: 10.3389/fonc.2018.00370.

Fura JM, Sabulski MJ, Pires MM. (2014). D-Amino Acid Mediated Recruitment of Endogenous Antibodies to Bacterial Surfaces. Acs Chemical Biology. 9(7):1480–9 doi 10.1021/cb5002685.

Galdiero MR, Garlanda C, Jaillon S, Marone G, Mantovani A. (2012). Tumor associated macrophages and neutrophils in tumor progression. J Cell Physiol. doi: 10.1002/jcp.24260

Garbe C, Krasagakis K, Zouboulis CC, Schröder K, Krüger S, Stadler R, Orfanos CE. (1990). Antitumor activities of interferon-alpha, interferon-beta, and interferon-gamma and their combinations on human-melanoma cells-invitro-changes of proliferation, melanin synthesis, and immunophenotype. Journal of Investigative Dermatology. 95(6):S231–S7 doi 10.1111/1523-1747.ep12875837.

Hou DY, Muller AJ, Sharma MD, DuHadaway J, Banerjee T, Johnson M, et al. (2007). Inhibition of indoleamine 2,3-dioxygenase in dendritic cells by stereoisomers of 1-methyl-tryptophan correlates with antitumor responses. Cancer Research. 67(2):792–801 doi 10.1158/0008-5472.can-06-2925.

Jehs T, Faber C, Juel HB, Bronkhorst IH, Jager MJ, Nissen MH, et al. (2014) Inflammation-Induced Chemokine Expression in Uveal Melanoma Cell Lines Stimulates Monocyte Chemotaxis. Investigative Ophthalmology & Visual Science. 55(8):5169–75 doi 10.1167/iovs.14-14394.

Liu KT, Liu YH, Liu HL, Chong IW, Yen MC, and Kuo PL. (2016). Neutrophils are Essential in Short Hairpin RNA of Indoleamine 2,3-Dioxygenase Mediated-antitumor Efficiency. Mol Ther Nucleic Acids. doi: 10.1038/mtna.2016.105.

Lollini PL, Bosco MC, Cavallo F, De Giovanni C, Giovarelli M, Landuzzi L, et al. (1993). Inhibition of tumor-growth and enhancement of metastasis after transfection of the gamma-interferon gene. International Journal of Cancer. 55(2):320–9 doi 10.1002/ijc.2910550224.

Mojic M, Takeda K, Hayakawa Y. (2017). The Dark Side of IFN-γ: Its Role in Promoting Cancer Immunoevasion. Int J Mol Sci. doi: 10.3390/ijms19010089.

Moreno AC, Clara RO, Coimbra JB, Júlio AR, Albuquerque RC, Oliveira EM, et al. (2013). The expanding roles of 1-methyl-tryptophan (1-MT): in addition to inhibiting kynurenine production, 1-MT activates the synthesis of melatonin in skin cells. Febs Journal. 280(19):4782–92 doi 10.1111/febs.12444.

Platten M, Doeberitz NK, Oezen I, Wick W and Ochs K. (2015). Cancer immunotherapy by targeting IDO1/TDO and their downstream effectors. Front. Immunol. doi: 10.3389/fimmu.2014.00673.

Prendergast GC, Smith C, Thomas S, Mandik-Nayak L, Laury-Kleintop L, Metz R, et al. (2014). Indoleamine 2,3-dioxygenase pathways of pathogenic inflammation and immune escape in cancer. Cancer Immunology Immunotherapy. 63(7):721–35. doi: 10.1007/s00262-014-1549-4.

Qian F, Liao J, Villella J, Edwards R, Kalinski P, Lele S, et al.(2012). Effects of 1-methyltryptophan stereoisomers on IDO2 enzyme activity and IDO2-mediated arrest of human T cell proliferation. Cancer Immunology Immunotherapy. 61(11):2013–20 doi: 10.1007/s00262-012-1265-x.

Radkov AD, Moe LA. (2014). Bacterial synthesis of D-amino acids. Applied Microbiology and Biotechnology. 98(12):5363–74. doi 10.1007/s00253-014-5726-3.

Schiller JH, Pugh M, Kirkwood JM, Karp D, Larson M, Borden E. (1996). Eastern cooperative group trial of interferon gamma in metastatic melanoma: An innovative study design. Clinical Cancer Research. 2(1):29–36.

Selvan, Senthamil R., Dowling, John P., Kelly, William K., et al. (2016). Indoleamine 2,3-dioxygenase (IDO): Biology and Target in Cancer Immunotherapies. Current Cancer Drug Targets. 16,9, 755–764.

Tang SC. Montero AJ, Munn D, Link C, Vahanian, Kennedy E, Solima H. (2016) A phase 2 randomized trial of the 100 pathway inhibitor indoximod in combination with taxane based chemotherapy for metastatic breast cancer: Preliminary data. Cancer Research. 76. doi: 10.1158/1538-7445.sabcs15-p2-11-09.

Voo KS, Bover L, Harline ML, Weng J, Sugimoto N, Liu YJ. (2014). Targeting of TLRs inhibits CD4+ regulatory T cell function and activates lymphocytes in human peripheral blood mononuclear cells. J Immunol. 193(2):627–34. doi: 10.4049/jimmunol.1203334.

Yentz S, Smith D. (2018). Indoleamine 2,3‑Dioxygenase (IDO) Inhibition as a Strategy to Augment Cancer Immunotherapy. BioDrugs. doi: 10.1007/s40259-018-0291-4.

Zaidi MR, Merlino G. (2011). The Two Faces of Interferon-gamma in Cancer. Clinical Cancer Research. 17(19):6118–24 doi: 10.1158/1078-0432.ccr-11-0482.

Zhai L, Spranger S, Binder DC, Gritsina G, Lauing KL, Giles FJ, Wainwright DA (2015). Molecular Pathways: Targeting IDO1 and Other Tryptophan Dioxygenases for Cancer Immunotherapy. Clinical Cancer Research. 21(24):5427–33. doi: 10.1158/1078-0432.CCR-15-0420.

